# Distinct autoinhibitory mechanisms regulate vinculin binding by αT-catenin and αE-catenin

**DOI:** 10.1101/2020.10.25.354415

**Authors:** Jonathon A. Heier, Sabine Pokutta, Ian W. Dale, Sun Kyung Kim, Andrew P. Hinck, William I. Weis, Adam V. Kwiatkowski

## Abstract

α-catenin binds directly to β-catenin and connects the cadherin-catenin complex to the actin cytoskeleton. Tension regulates α-catenin conformation: actomyosin-generated force stretches the middle(M)-region to relieve autoinhibition and reveal a binding site for the actin-binding protein vinculin. Here we describe the biochemical properties of αT(testes)-catenin, an α-catenin isoform critical for cardiac function, and how intramolecular interactions regulate vinculin binding autoinhibition. Isothermal titration calorimetry (ITC) showed that αT-catenin binds the β-catenin/N-cadherin complex with a similar low nanomolar affinity to that of αE-catenin. Limited proteolysis revealed that the αT-catenin M-region adopts a more open conformation than αE-catenin. The αT-catenin M-region binds the vinculin N-terminus with low nanomolar affinity, indicating that the isolated αT-catenin M-region is not autoinhibited and thereby distinct from αE-catenin. However, the αT-catenin head (N- and M-regions) binds vinculin 1000-fold more weakly (low micromolar affinity), indicating that the N-terminus regulates M-region binding to vinculin. In cells, αT-catenin recruitment of vinculin to cell-cell contacts requires the actin-binding domain and actomyosin-generated tension, indicating that force regulates vinculin binding. Together, our results indicate that the αT-catenin N-terminus is required to maintain M-region autoinhibition and modulate vinculin binding. We postulate that the unique molecular properties of αT-catenin allow it to function as a scaffold for building specific adhesion complexes.

## Introduction

The cadherin-catenin-complex that forms the core of the adherens junction (AJ) is required for intercellular adhesion and tissue integrity (1–3). Classical cadherins are single-pass transmembrane proteins with an extracellular domain that forms *trans* interactions with cadherins on adjacent cells (4–6). The adhesive properties of classical cadherins are driven by the recruitment of cytosolic catenin proteins to the cadherin tail: p120-catenin binds to the juxta-membrane domain and β-catenin binds to the distal part of the tail. β-Catenin recruits α-catenin, a mechanoresponsive actin-binding protein (7–13). The AJ mechanically couples and integrates the actin cytoskeletons between cells to allow dynamic adhesion and tissue morphogenesis (3).

The best characterized member of the α-catenin family of proteins is mammalian αE(epithelial)-catenin. Structurally, it is composed of 5 four-helix bundles connected to a C-terminal five-helix bundle by a flexible linker (14–16). The two N-terminal four-helix bundles form the N-domain that binds β-catenin and mediates homodimerization (12,17–19). The middle three four-helix bundles form the middle (M)-region that functions as a mechanosensor (20–25). The C-terminal five-helix bundle forms the F-actin binding domain (ABD) (11,13,26,27). F-actin binding is allosterically regulated. αE-catenin can bind F-actin readily as a homodimer, but when in complex with β-catenin, mechanical force is required for strong F-actin binding (9–11,26). In addition, when tension is applied to αE-catenin, salt bridge interactions within the M-region are broken, allowing the domain to unfurl to reveal cryptic binding sites for other cytoskeletal binding proteins such as vinculin (16,23–25,28–32). The recruitment of these proteins is thought to help stabilize the AJ in response to increased tension and further integrate the F-actin cytoskeleton across cell-cell contacts (24,28,31,33–35).

Three α-catenin family proteins are expressed in mammals: the ubiquitous αE-catenin, αN (Neuronal)-catenin and αT (Testes)-catenin (36,37). αE-catenin and αN-catenin are 81% identical and 91% similar in amino acid sequence. αT-catenin is 58-59% identical and 77% similar to αE-catenin and αN-catenin, making it the most divergent of the family(36–38). αT-catenin is predominantly expressed in the heart and testes (39). In the heart, it localizes to the intercalated disc (ICD), a specialized adhesive structure that maintains mechanical coupling and chemical communication between cardiomyocytes (40,41). In mice, loss of αT-catenin from the heart causes dilated cardiomyopathy and mutations in αT-catenin are linked to arrhythmogenic ventricular cardiomyopathy in humans (42,43). In addition to cardiomyopathy, αT-catenin is linked to multiple human diseases, including asthma, neurological disease and cancer (44–46).

Despite a growing realization of its importance in human disease, the molecular properties and ligand interactions of αT-catenin remain poorly understood. Our previous work revealed that αT-catenin, unlike mammalian αE-catenin, is a monomer in solution that can bind to F-actin with low micromolar affinity in the absence of tension. F-actin binding is also not allosterically regulated, as the β-catenin/αT-catenin complex binds to F-actin with the same affinity as αT-catenin monomer (38). Single molecule pulling experiments have shown the αT-catenin M-region to be mechanoresponsive as it unfurls in a force range similar to αE-catenin (47).

Here we show that αT-catenin associates with the components of the cadherin/catenin complex with the same affinity as αE-catenin *in vitro*, revealing that they may compete with one another for binding at the plasma membrane. We also show that the M-region of αT-catenin is not autoinhibited and can bind the vinculin N-terminus in the absence of tension with strong affinity. This result indicates a distinct method of regulation when compared to the tension dependent mechanism of the αE-catenin M-region. However, when the N-terminus of αT-catenin is attached to the M-region, we found the affinity for the N-terminus of vinculin drops significantly. This indicates that interdomain interactions between the N-terminus and the M-region of αT-catenin regulate its interaction with vinculin. We measured the recruitment of vinculin to cell-cell contacts and found that despite the distinct mechanism of regulation, recruitment of vinculin is still tension dependent. We postulate that this mechanism is important for the ability of αT-catenin to build AJs in a manner distinct from that of αE-catenin.

## Results

### αT-catenin binds tightly to the β-catenin/N-cadherin core complex

We characterized the interaction between αT-catenin and β-catenin by isothermal titration calorimetry (ITC) using purified recombinant proteins. We used the head region (comprising the N- and M-domains) of αT-catenin (aa1-659, Fig. 1A) for these experiments since it is more stable than fulllength αT-catenin and yields sufficiently high protein concentrations for ITC. Past studies revealed that αE-catenin head region (aa1-651) binds β-catenin and the β-catenin•E-cadherin tail complex with a similar affinity to full-length αE-catenin (12). We observed that the αT-catenin head binds β-catenin with a dissociation constant ~250 nM (Fig. 1B; Table 1). The affinity of αT-catenin for β-catenin is an order of magnitude weaker than the association of αE-catenin or αN-catenin for β-catenin (15-20 nM; (12).

**Figure 1.**
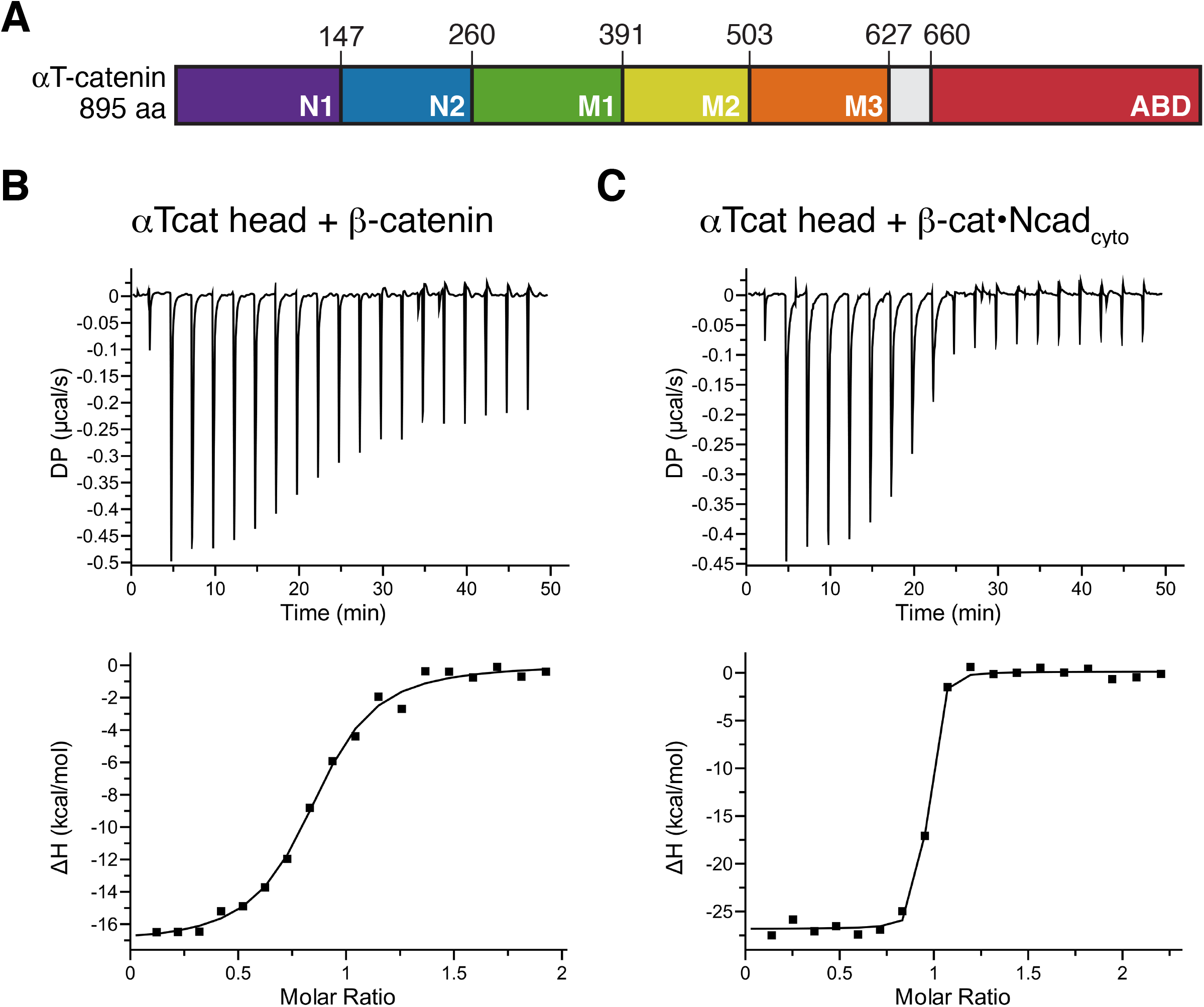
αT-catenin binds the N-cadherin•β-catenin complex with nanomolar affinity. A. Domain organization of αT-catenin. Amino acid domain boundaries marked. B-C. αT-catenin head region (aa1-659, αTcat head) binding to β-catenin (B) or the β-catenin•N-cadherin cytoplasmic tail (β-cat•Ncad_cyto_) complex (C) was measured by ITC. The ratio of heat released (kcal) per mole of β-catenin or β-cat•Ncad_cyto_ injected into αT-catenin head was plotted against the molar ratio of αT-catenin head and β-catenin or αT-catenin head and β-cat•Ncad_cyto_. Thermodynamic properties derived from these traces are shown in Table 1.

**Table 1.** ITC measurements of αT-catenin fragments binding to β-catenin or β-catenin•N-cadherin cytoplasmic tail complex.

| Proteins | $K_d$<br>(nM) | $\Delta H$<br>(kcal/mol) | $T\Delta S$<br>(kcal/mol) | $\Delta G$<br>(kcal/mol) | # |
| --- | --- | --- | --- | --- | --- |
| αT-catenin head (1-659) + β-catenin | $264.1 \pm 109.1$ | $-14.6 \pm 2.1$ | -5.6 | -9.0 | 5 |
| αT-catenin head (1-659) + β-catenin•Ncad<sub>cyto</sub> | $5.6 \pm 0.6$ | $-27.0 \pm 0.2$ | -15.8 | -11.3 | 3 |
| αT-catenin N1-N2 (1-259) + β-catenin•Ncad<sub>cyto</sub> | $6.9 \pm 12.0$ | $-18.9 \pm 1.0$ | -7.8 | -11.1 | 1 |

Cadherin tail binding to β-catenin strengthens the affinity between β-catenin and α-catenin (12). N-cadherin is the primary classical cadherin expressed in cardiomyocytes (48). We tested if the N-cadherin tail (Ncad_cyto_) influences the αT-catenin/β-catenin interaction by titrating β-catenin•Ncad_cyto_ complex into αT-catenin head (Fig. 1C). The affinity of this interaction was 5-6 nM (Table 1), indicating that αT-catenin binds to the cadherin•β-catenin complex an order of magnitude more strongly than to β-catenin alone. This affinity is similar to the 1-2 nM affinity observed between the cadherin tail•β-catenin complex and αE-catenin or αN-catenin (12) and suggests that αT-catenin can effectively compete with αE-catenin for binding to the membrane-associated cadherin•β-catenin complex.

### αT-catenin N-terminus is monomeric

Full-length αT-catenin is primarily a monomer in solution, though it does have homodimerization potential *in vitro* (38). The best evidence for dimerization potential comes from a point mutation linked to arrhythmogenic ventricular cardiomyopathy in humans, V94D, that renders αT-catenin an obligate homodimer (38,43). We analyzed the oligomerization properties of the αT-catenin N1-N2 (aa1-259, Fig. 1A) and compared them to the V94D mutant. Analytical size exclusion chromatography (SEC) of αT-catenin wild type (wt) N1-N2 and V94D N1-N2 revealed that wt N1-N2 eluted as a single monomer species (Fig. 2A, blue line) whereas the V94D mutant eluted as a dimer species (Fig 2A, red line).

**Figure 2.**
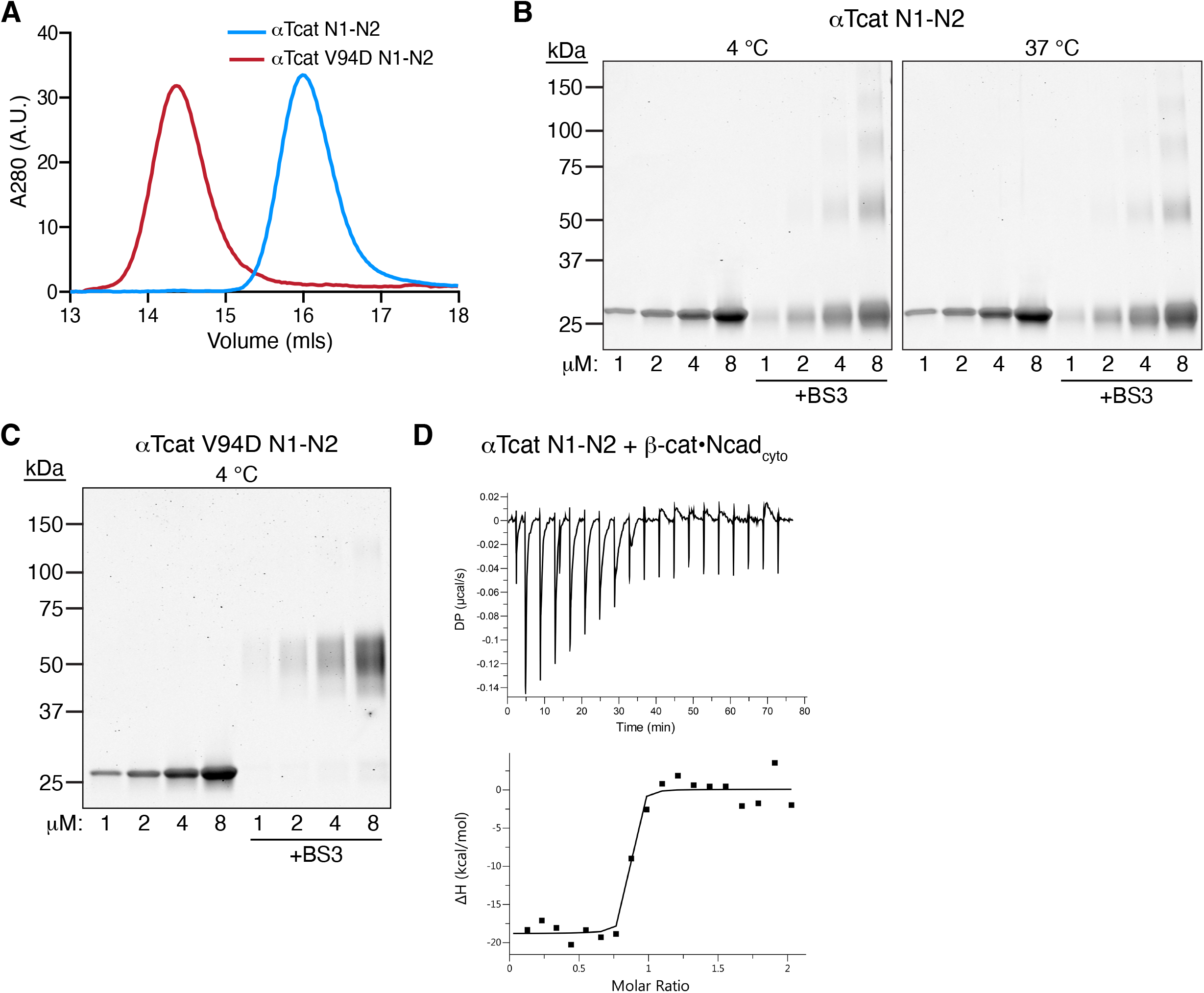
αT-catenin N1-N2 is a monomer in solution. A. Analytical SEC of 30 μM αT-catenin N1-N2 (aa1-259, blue line) and 30 μM αT-catenin V94D N1-N2 (red line). B. Crosslinking experiments with αT-catenin N1-N2. Increasing concentrations of αT-catenin a were incubated with or without 1 mM BST at 4 °C (left panel) or 37 °C (right panel) for 30 minutes, separated by SDS-PAGE and stained with Coomassie dye. C. Crosslinking experiments with αT-catenin V94D N1-N2. Increasing concentrations of αT-catenin V94D N1-N2 were incubated with or without 1 mM BST at 4 °C for 30 minutes, separated by SDS-PAGE and stained with Coomassie dye. D. αT-catenin N1-N2 binding to β-catenin•N-cadherin cytoplasmic tail (β-cat•Ncad_cyto_) was measured by ITC. Thermodynamic properties derived from this trace are shown in Table 1.

We then analyzed the oligomeric state of the αT-catenin N-terminus by cross-linking. Increasing concentrations of αT-catenin N1-N2 were incubated with or without the cross-linker bis(sulfosuccinimidyl)suberate (BS3) at 4 °C or 37 °C and the resulting products analyzed by SDS-PAGE. As expected, the αT-catenin N1-N2 migrated as a 25 kDa protein in the absence of cross-linker (Fig. 2B). Incubation with BS3 did not affect migration at low concentrations, although at higher concentrations (4 and 8 μM), larger species were detected at both temperatures. In contrast, αT-catenin V94D N1-N2 ran as 50 kDa protein in the presence of BS3 at all concentrations tested (Fig. 2C), indicating a cross-linked dimer. We conclude that the αT-catenin N-terminus, similar to full-length protein, is primarily a monomer in solution.

We then tested the ability of the αT-catenin N1-N2 to bind to the β-catenin•Ncad_cyto_ complex by ITC. The affinity of this interaction was 7 nM (Fig. 2D, Table 1), similar to the αT-catenin head and confirming that this fragment contains the complete β-catenin binding site.

### αT-catenin M-region binds vinculin

The αE-catenin M-region is autoinhibited with respect to vinculin binding: mechanical force is required to break an internal salt bridge network within M1-M3 to reveal the vinculin binding site in M1 (25,30). The αT-catenin M-region also recruits vinculin and force is required to unfurl M1 to promote high affinity binding (47). However, our previous proteolysis experiments of full-length αT-catenin revealed that the M2M3 region existed in a more open, protease-sensitive state (38). Notably, the amino acids that form the salt bridges required for autoinhibition in the αE-catenin M-region are conserved in αT-catenin, with the exception of E277 in M1 that pairs with R451 in M2. In αT-catenin, the arginine is conserved at the corresponding residue (aa446), but the glutamic acid is replaced by a threonine at aa272 (25), preventing these residues from interacting. We questioned if αT-catenin M1-M3 adopted an autoinhibited conformation.

We examined the organization and vinculin binding properties of the complete αT-catenin M-region (M1-M3, aa260-626). αT-catenin M1-M3 eluted as a single, discrete peak by SEC, identical to αE-catenin M1-M3 (aa273-651; Fig. 3A). We then probed the M-region flexibility by limited trypsin proteolysis. Tryptic digestion of αE-catenin M1-M3 revealed that it was largely protease-resistant: nearly 50% of the fragment was still intact after 120 minutes of digestion, consistent with it forming a closed, autoinhibited domain (Fig. 3B). In contrast, αT-catenin M1-M3 was completely digested after 60 minutes into three stable fragments at 23, 16 and 12 kDa (Fig. 3B). Note that both M-regions contain a similar number of lysine and arginine residues (47 in αE-catenin and 39 in αT-catenin), the majority of which are conserved. N-terminal sequencing revealed that the 23 kDa and 16 kDa fragments both started at aa379 and represented the M2-M3 and M2 bundles, respectively. The 12 kDa fragment started at aa485 and corresponded to the M3 bundle (Fig. 3C). Similar protease resistant fragments were observed from digest of full-length protein (38) and are consistent with the αT-catenin M-region adopting a more open, protease-sensitive state relative to αE-catenin, despite five of the six salt bridge residue pairs being conserved. Likewise, the isolated αT-catenin M1-M3 fragment does not appear to adopt a stable, autoinhibited conformation.

**Figure 3.**
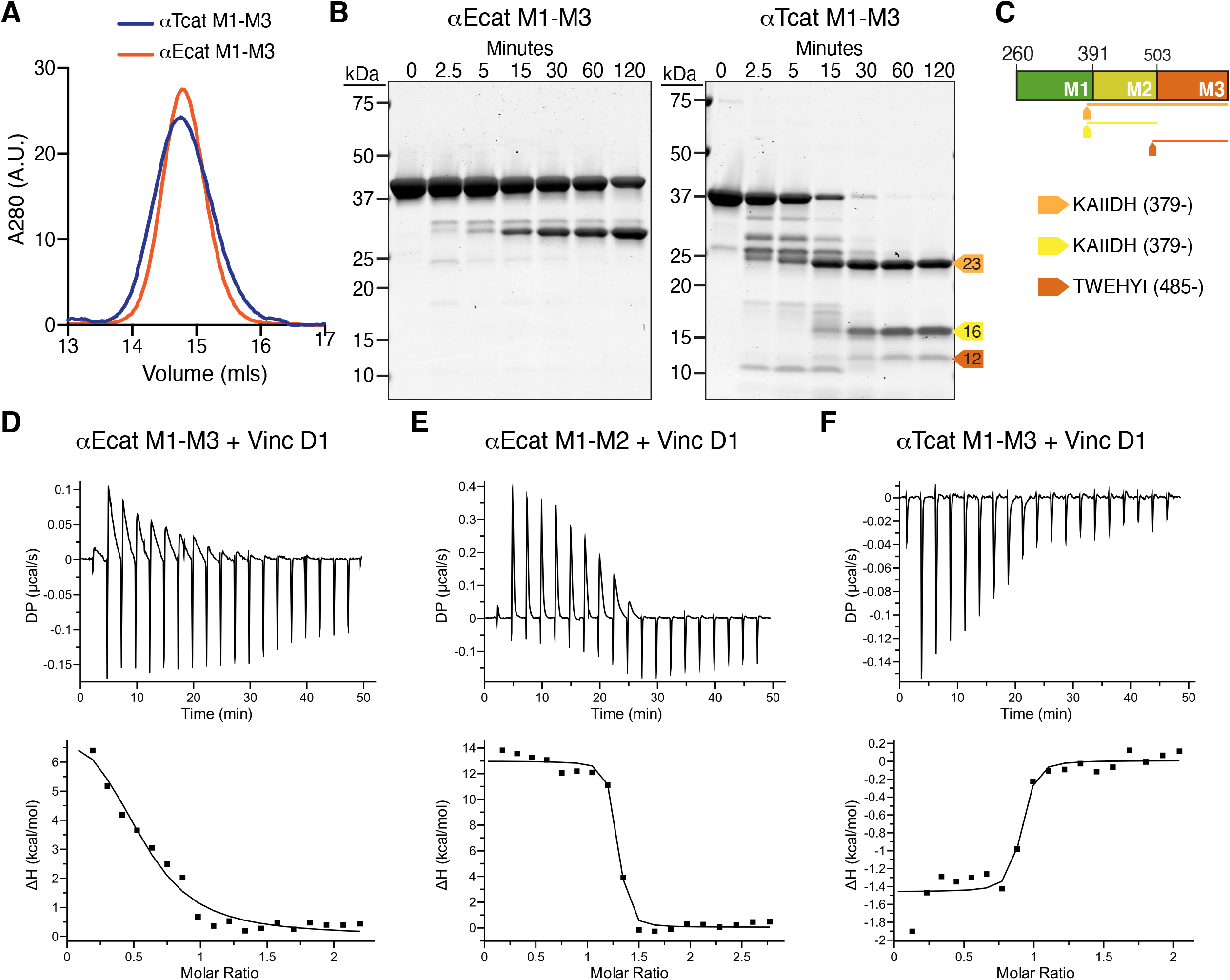
αT-catenin M1-M3 binds vinculin D1 with high affinity. A. Analytical SEC of recombinant αT-catenin M1-M3 (aa260-626) and αE-catenin M1-M3 (aa273-651). B. Limited proteolysis of αT-catenin and αE-catenin M1-M3 fragments. Proteins incubated for 0, 2.5, 5, 15, 30, 60, and 120 min at room temperature in 0.05 mg/ml trypsin, resolved by SDS-PAGE and stained with Coomassie dye. Stable fragments of 23 (yellow-orange), 16 (yellow) and 12 kDa (orange) are marked with colored arrows. C. Edman sequencing results of limited proteolysis fragments. Protein fragments are mapped on the M1-M3 sequence as color-coded lines. D-F. Representative ITC traces of αE-catenin M1-M3 (D), αE-catenin M1-M2 (aa273-510, E) and αT-catenin M1-M3 (F) binding to vinculin D1 (aa1-259). Thermodynamic properties derived from these traces are shown in Table 2.

We then measured the affinity of αT-catenin M1-M3 for vinculin. We used the vinculin D1fragment (aa1-259) which contains the first two four-helix bundles and binds to αE-catenin with a similar affinity to the vinculin head domain, D1-D4 (23). As previously observed (23), the autoinhibited αE-catenin M1-M3 fragment bound weakly to vinculin D1 (Fig. 3D, Table 2). When M3 was deleted from this fragment, auto-inhibition was relieved and αE-catenin M1-M2 bound to vinculin with low nanomolar affinity, as observed previously (Fig. 3E, Table 2) (23). In contrast, the αT-catenin M1-M3 fragment showed strong, nanomolar binding to vinculin D1 similar to αE-catenin M1-M2 (Fig. 3F, Table 2), indicating that αT-catenin M1-M3 binding to vinculin was not autoinhibited. Whereas binding to αE-catenin M1-M2 or M1-M3 is endothermic (entropy driven), consistent with unfolding of the M1 bundle needed for this interaction (23), binding to αT-catenin M1-M3 was exothermic, suggesting that αT-catenin M1 is unfurled in the M1-M3 fragment.

**Table 2.**
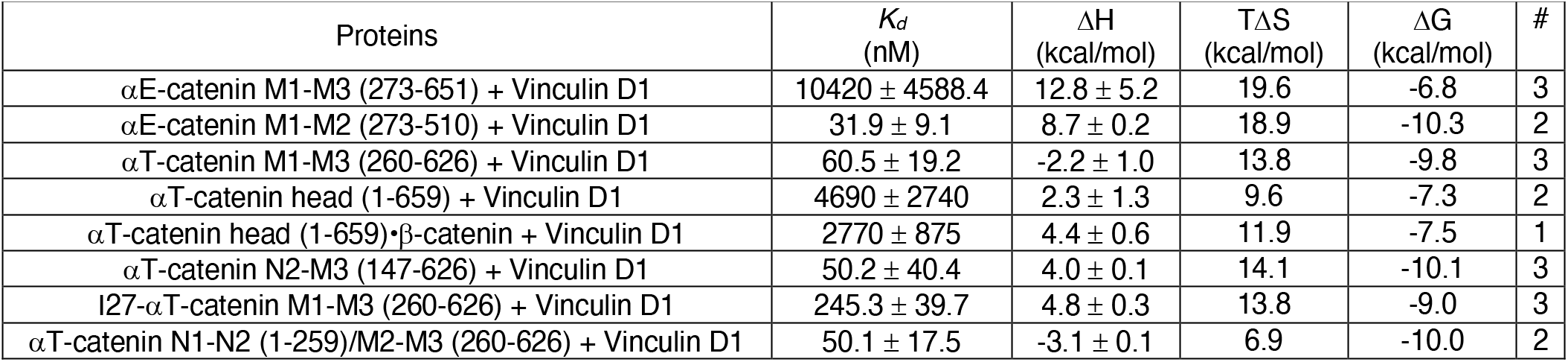
ITC measurements of αT-catenin and αE-catenin M-domain fragments binding to vinculin D1.

### αT-catenin N-terminus regulates vinculin binding

Recent *in vitro* single molecule stretching experiments revealed that force is required to expose the vinculin binding site in αT-catenin M1-M3 (47). However, our ITC results with the αT-catenin M1-M3 fragment indicated that tension was not required to release M1. In the stretching experiments, the αT-catenin M1-M3 fragment was flanked by a pair of titin I27 immunoglobulin-like domains and the N-terminus of the fusion protein was tethered to a substrate (47). We questioned if N-terminal associations with M1-M3 stabilize M1 and regulate vinculin binding.

We first characterized the interaction between αT-catenin head domain and vinculin D1 by ITC. The binding shifted from exothermic to endothermic and the affinity was ~5 μM, two orders of magnitude weaker than observed with M1-M3 (Fig. 4A, Table 2). This suggested that the addition of N1-N2 stabilized M1 and inhibited vinculin binding.

**Figure 4.**
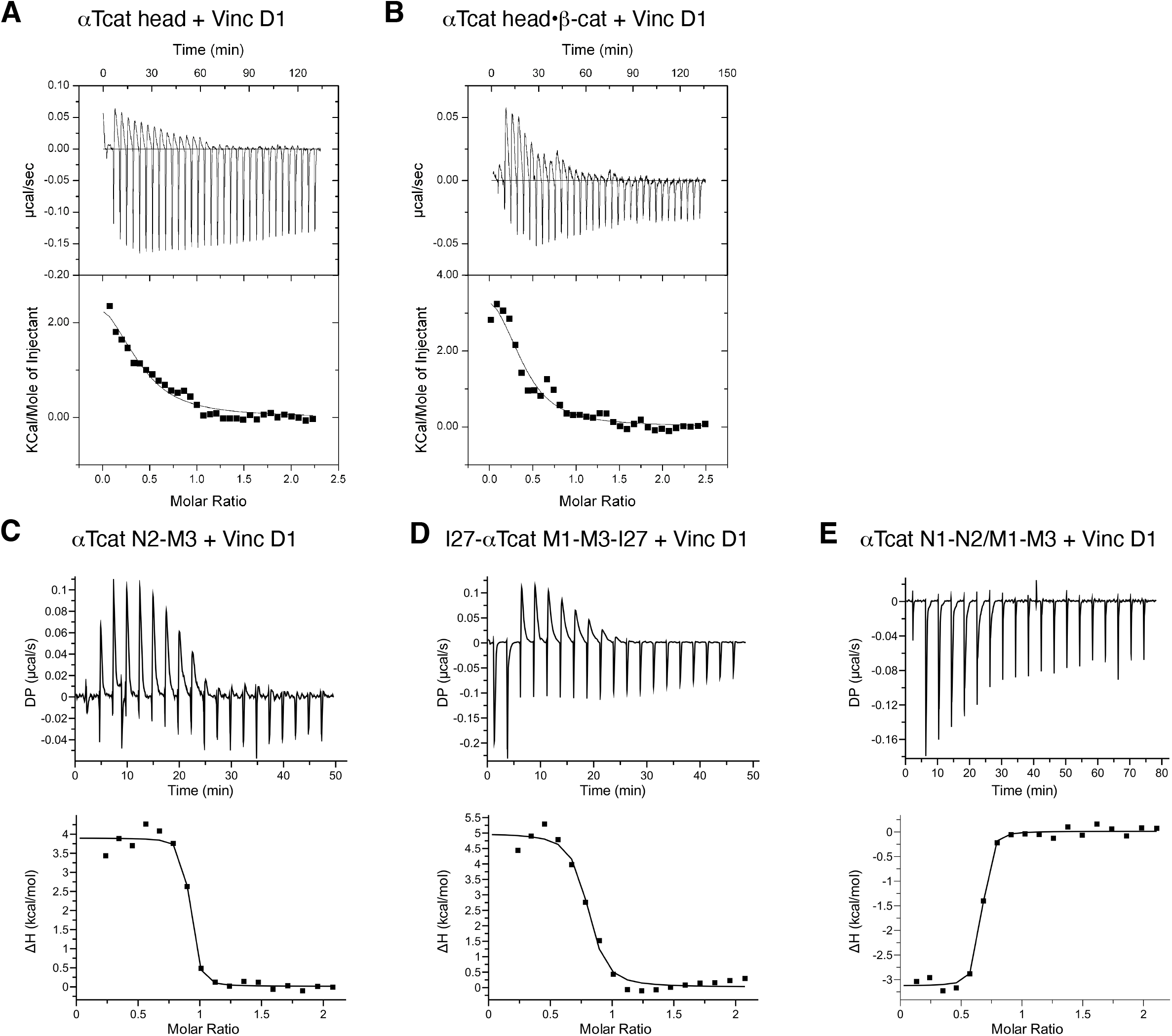
αT-catenin N-terminus regulates vinculin binding to M1-M3. A-D. Representative ITC traces of vinculin D1 binding to αT-catenin head (A), αT-catenin head/β-catenin complex, αT-catenin N2-M3 (aa147-626), 2I27-αT-catenin M1-M3(aa259-667)-2I27 (D), and αT-catenin N1-N2/M1-M3 complex (E). Thermodynamic properties derived from these traces are shown in Table 2.

Recent work revealed allosteric coupling between the N-terminal domains and M-region of αE-catenin (16). Specifically, the presence of β-catenin caused changes in the accessibility of cysteine residues in the N2-M2 interface and in M3. Since β-catenin binding alters the relative positions of the αE-catenin N1 and N2 domains (12), we tested if β-catenin binding to the αT-catenin N-region affected vinculin binding to the M-region. Vinculin D1 was titrated into a solution of purified αT-catenin head•β-catenin complex. The presence of β-catenin had little impact on the affinity (~3 μM, Fig. 4B, Table 2), indicating that N-terminal-mediated autoinhibition was maintained. This result is consistent with past work showing that the β-catenin•cadherin complex had no effect on αE-catenin binding to vinculin (23).

We then asked if the entire N-terminus is required to regulate M1 binding. The addition of N2 to αT-catenin M1-M3 (aa147-626) did not weaken vinculin D1 binding relative to M1-M3 (*K_d_* = 50 nM, Fig. 4C, Table 2). The reaction switched from exothermic to endothermic, suggesting partial compensation of M1 stability in this fragment. Thus, all or part of N1 is required to regulate M1 interactions with vinculin. This is consistent with the observation that removing the first 56 residues of N1 from full-length αE-catenin reduces the inhibition of vinculin binding by about 50 fold (23).

We next questioned if the titin repeats attached to M1-M3 in the construct used by Pang, Le and colleagues (47) stabilized M1. We titrated vinculin D1 into a solution of the 2I27-αT-catenin M1-M3(aa259-667)-2I27 construct. Notably, the binding was endothermic and the affinity was ~250 nM, markedly weaker than observed with αT-catenin M1-M3 alone (Fig. 4D, Table 2). We speculate that the well-structured titin repeats promote M1 stability and may sterically occlude vinculin D1 binding, thus reducing the affinity. The M1 stability and steric hindrance provided by the I27 repeats partially mimic the complete N-terminus and likely explain why tension is needed to promote vinculin binding in the single molecule stretching experiments whereas αT-catenin M1-M3 in solution binds vinculin readily.

Given that the entire N-terminus is required to regulate vinculin binding in αT-catenin, we asked if N1-N2 could regulate M1-M3 in *trans*. We titrated vinculin D1 into a equimolar mixture of αT-catenin N1-N2 and M1-M3. The binding reaction was similar to M1-M3 alone: exothermic with a *K_d_* of 50 nM (Fig. 4E, Table 2). This result indicates that the M-region must be physically coupled to the N-terminus to stabilize M1 and regulate ligand accessibility.

### Tension recruits vinculin to αT-catenin

We tested the ability of αT-catenin to restore cellcell adhesion and recruit vinculin in α-catenin-deficient R2/7 carcinoma cells (49). We transiently expressed EGFP-αE-catenin or EGFP-αT-catenin in R2/7 cells and analyzed cell-cell contact formation and endogenous vinculin recruitment by immunostaining. Both EGFP-αE-catenin and EGFP-αT-catenin restored cell-cell adhesion, organized F-actin along cell-cell contacts and recruited vinculin to junctions (Fig. 5). To determine if vinculin recruitment was tension dependent, we treated cells with DMSO or 50 μM blebbistatin to suppress myosin activity for 30 minutes and stained for vinculin (Fig. 5, A-D). αE-catenin and αT-catenin recruited similar levels of vinculin in DMSO controls (Fig. 5, A, B and E). Importantly, blebbistatin treatment significantly reduced vinculin recruitment to both αE-catenin and αT-catenin containing AJs (Fig. 5, C, D and E). Thus, myosin-based tension is required to recruit vinculin to αT-catenin, similar to αE-catenin.

**Figure 5.**
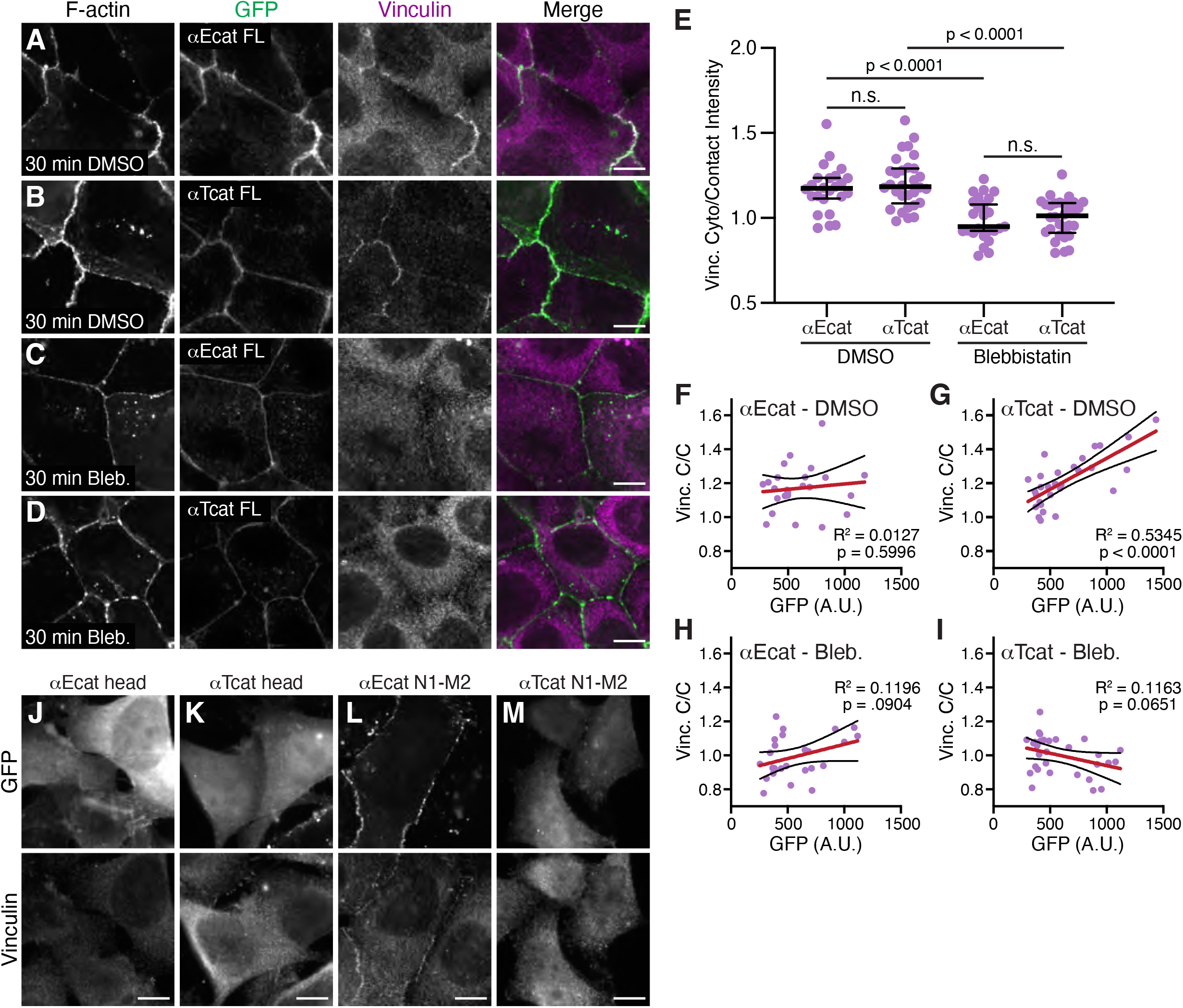
αT-catenin recruitment of vinculin to cell-cell contacts is tension-dependent. A-D. R2/7 cells were transfected with EGFP-αE-catenin full length (aEcat FL; A and C) or EGFP-αT-catenin full length (αTcat FL; B and D) and treated for 30 mins with DMSO (A and B) or 50 μM blebbistatin (C and D) before fixation. Cells were stained for F-actin and vinculin. Individual and merged vinculin (magenta) and GFP (green) channels shown. E. Quantification of vinculin intensity at cell-cell contacts. Signal intensity at contacts was divided by the average cytoplasmic intensity and a scatter plot of all data points is shown. The black horizontal line is the median and the error bars define the interquartile range. Oneway ANOVA with Tukey’s comparisons; n.s. = not significant. n ≥ 24 images from two independent experiments. F-I. Vinculin contact/cytoplasmic ratio (C/C) was plotted against the average GFP intensity of each fusion construct from the masked cell contact region. Linear regression analysis was performed to calculate the slope (red line), 95% confidence intervals (black lines surrounding slope) and R^2^ value. The slope deviation from zero was analyzed for significance (p value). n ≥ 24 images from two independent experiments. J-M. R2/7 cells were transfected with EGFP-tagged αE-catenin head (aa1-670, J), αT-catenin head (aa1-659, K), αE-catenin N1-M2 (aa1-510, L) and αT-catenin N1-M2 (aa1-502, M), fixed and stained for vinculin. Individual GFP (top panel) and vinculin (bottom panel) channels shown. Scale bar is 10 μm in all images.

We then examined the correlation between EGFP fusion expression and vinculin recruitment. As expected, EGFP-αE-catenin expression levels did not correlate with vinculin levels since vinculin was only recruited when αE-catenin was activated by sufficient tension (24,28). However, we observed a strong positive correlation between EGFP-αT-catenin expression and vinculin recruitment. We observed a similar positive correlation with constitutively-open αE-catenin constructs in cardiomyocytes (34). Together, these results suggest that while vinculin recruitment to αT-catenin is tension dependent, the force threshold for binding may be low, permitting all/most AJ-incorporated αT-catenin molecules to recruit vinculin.

Finally, we tested if actin binding was required to relieve αT-catenin autoinhibition and if removal of M3 opened the M-region to permit vinculin recruitment. We expressed EGFP-tagged truncations of αE-catenin and αT-catenin: the head domain lacking the ABD (N1-M3) or the head domain minus M3 and ABD (N1-M2). As expected, αE-catenin and αT-catenin N1-M3 were both cytosolic and unable to form cell-cell contacts and recruit vinculin (Fig. 5, J and K). Deletion of M3 in αE-catenin relieves autoinhibition, and αE-catenin N1-M2 (aa1-510) was able to recruit vinculin and restore cell-cell adhesion, as shown previously (49). Here vinculin provides the necessary actin binding activity to link the cadherin-catenin complex to actin through αE-catenin. In contrast, αT-catenin N1-M2 (aa1-502) was cytosolic and failed to recruit vinculin to cellcell contacts. This suggests that, in αT-catenin, loss of M3 does not release M1 for ligand binding and underscores the importance of the N-terminus in regulating M1 binding events.

## Discussion

### αT-catenin forms a strong AJ core with cadherin•β-catenin

Our ITC results show that the β-catenin•N-cadherin tail complex binds with high, ~5 nM affinity to αT-catenin similar to 1-2 nM affinity previously reported between the β-catenin•cadherin tail complex and αE-catenin (12). Thus αT-catenin forms a strong cadherin-catenin core complex like αE-catenin to link actin to the AJ. αT-catenin is coexpressed with αE-catenin in multiple mammalian tissues and, in the heart, it is enriched along the ICD with αE-catenin (39,44,45,50). It is unknown if αE-catenin and αT-catenin bind stochastically to the cadherin•β-catenin complex in cardiomyocytes or if binding is regulated to favor one α-catenin or the other. Notably, loss of either αE-catenin or αT-catenin from the mouse heart causes dilated cardiomyopathy (42,51), suggesting that each α-catenin has a unique, critical role in ICD function and heart physiology. In addition, αE-catenin and αT-catenin (Heier and Kwiatkowski, unpublished data) also bind plakoglobin, the desmosome-associated β-catenin homolog that is enriched at the ICD (52,53). Plakoglobin binds the cadherin tail with an affinity similar to β-catenin and both plakoglobin and β-catenin are enriched at cardiomyocyte AJs (50). The downstream consequences of α-catenin binding to plakoglobin or β-catenin are not clear, but recent evidence that the N-terminus and M-region of αE-catenin are allosterically coupled (16) raises the possibility that differential binding could affect downstream ligand recruitment. Future work is expected to reveal how specific cadherin•β-catenin/plakoglobin•α-catenin complexes are formed and function to regulate adhesion and signaling.

In the absence of the cadherin tail, β-catenin alone binds αT-catenin an order of magnitude weaker than αE-catenin (~250 nM versus ~20 nM). This suggest that αT-catenin does not compete with αE-catenin for binding cytosolic β-catenin. Though there is evidence of a cytosolic αE-catenin•β-catenin complex in epithelial cells (54), the physiological relevance of this interaction is not clear and a similar α-catenin•β-catenin complex in cardiomyocytes has not been reported.

αE-catenin can homodimerize and the cadherin-free, cytosolic homodimer pool regulates cell actin dynamics and cell motility (9,10,12,54,55). In contrast, full-length αT-catenin is a monomer in solution and has limited dimerization potential with no evidence of homodimerization *in vivo* (38). Our results here show that the isolated αT-catenin N-terminus is also monomeric. We speculate that limited dimerization potential and weaker affinity for β-catenin favor a cadherin-free, cytosolic pool of αT-catenin monomer. The role of such a monomer pool is unclear.

### The αT-catenin M-region does not adopt an autoinhibited conformation in isolation

Structural, biochemical and biophysical data indicate that the αE-catenin M-region adopts an autoinhibited conformation (16,23–25,28–32). The vinculin binding site is buried in the folded M1 domain and interactions between M1, M2 and M3 maintain the autoinhibited form. Tension breaks these interactions to release M1, allowing it to unfold and bind strongly to vinculin. More recent biophysical data suggest that the αT-catenin M-region adopts a similar conformation with force being required to free M1 for high affinity binding to vinculin (47).

Our limited proteolysis experiments with the isolated αT-catenin M-region indicate that it exists in a more open, protease-sensitive conformation. Consistent with this, ITC results revealed that αT-catenin binds vinculin with low nanomolar affinity and is not autoinhibited. αT-catenin possesses 5 of the 6 residue pairs that form the salt bridge network that mediates αE-catenin autoinhibition (25). The 277E-451R bridge between M1 and M2 in αE-catenin is not conserved in αT-catenin, with the glutamic acid replaced by a threonine at aa272. The threonine at aa272 is conserved across the αT-catenin family. Assuming no major structural differences between the αE-catenin and αT-catenin M-regions, the 272T and 446R residues would not be able to interact. In simulations, the αE-catenin 277E-451R salt bridge is predicted to rupture quickly under external force (16,25). Unfortunately, we lack a structure of the αT-catenin M-region to determine if a similar network regulates domain organization. However, our data demonstrate that such intradomain interactions within the αT-catenin M-region are insufficient for autoinhibition and that the molecular requirements for autoinhibition differ between the two proteins.

### The N-terminus is required for αT-catenin M-region autoinhibition

The αT-catenin head fragment (N1-M3) showed weak, micromolar binding to vinculin, indicating that the N-terminus regulates M-region autoinhibition. The entire N-terminus (N1-N2) is required for autoinhibition since deleting N1 restored strong vinculin binding. Notably, replacing the N-terminus with another well-structured protein moiety, the titin I27 repeats, reduced vinculin binding and caused the binding reaction to switch from exothermic to endothermic, suggesting the M1 was stabilized. This observation explains the force-dependent vinculin binding to the αT-catenin M-region observed in recent biophysical experiments (47). The ability of the titin I27 repeats to partially restore autoinhibition also suggests that steric, nonspecific interactions rather than specific interdomain associations (e.g., salt bridges) between the αT-catenin N-terminus and M-region promote M1 folding and M-region autoinhibition.

### αT-catenin recruits vinculin

Expression of EGFP-αT-catenin was sufficient to restore cellcell contacts and recruit vinculin in α-catenin deficient R2/7 cells. Vinculin binding to αT-catenin requires actin binding by the latter and is tension-dependent, similar to αE-catenin. This is consistent with the vinculin binding site in αT-catenin M1 being occluded in the absence of force. Notably, deletion of M3 in αT-catenin did not relieve autoinhibition, offering additional evidence that the mechanism of autoinhibition is distinct from αE-catenin.

Vinculin is recruited to αE-catenin to bolster the AJ connection to actin under mechanical load (24,33,34,56). Neither the physiological context nor the functional role of vinculin recruitment to αT-catenin has been established *in vivo*. Loss of αE-catenin from the mouse heart disrupts AJ formation and causes a marked decrease in vinculin expression and recruitment, despite the presence of αT-catenin, resulting in cardiomyopathy (51). The inability of αT-catenin to compensate for αE-catenin in the heart underscores how the two α-catenins, despite sharing core properties, are regulated by distinct mechanisms and likely play unique roles in junction organization and signaling.

Together, our data support a model in which the αT-catenin N-terminus functions to regulate M-region stability and autoinhibition. Our *in situ* data indicate that force is required to reveal the vinculin binding site. We speculate that it may do so by separating the M-region from the N-terminus rather than breaking a network of internal M-region salt bridges. This could provide cadherin/β-catenin/αT-catenin complexes in cardiomyocytes with distinct mechanical properties, allowing ligand binding and allosteric signaling over a unique force scale relative to αE-catenin-containing complexes. While we have focused on vinculin binding to M1, intrinsic differences in αT-catenin M-region organization could also affect other M1 as well as M2 and M3 ligand interactions, possibly independent of force. Intramolecular and allosteric interactions are emerging as an important factor in the regulation of α-catenin conformation and molecular complex formation at the AJ. Further work is needed to define how molecular differences between the α-catenin protein family regulate mechanical adhesion and signaling.

## Experimental procedures

### Plasmids

Full-length *M. musculus* β-catenin, αT-catenin and αE-catenin as well as αT-catenin head region (aa1-659) in pGEX-TEV were described previously (9,12,38). The vinculin D1 construct (aa1-259) in pGEX-TEV was described in (23). *M. musculus* αT-catenin N1-N2 (aa1-259) N1-M3 (aa 1-659), M1-M3 (aa 260-626) N2-M3 (aa 147-626) and αE-catenin M1-M3 (aa273-651), M1-M2 (aa273-510) were cloned into pGEX-TEV. The constructs encoding αT-catenin M1-M3 flanked by I27 handles in pET151/D-TOPO was kindly provided to us by Dr. Jie Yan (47).

For mammalian cell expression, *M. musculus* αE-catenin fragments aa1-670 and aa1-510 and αT-catenin fragments aa1-659 and aa1-502 were cloned into pEGFP-C1.

### Recombinant Protein Expression and Purification

GST-tagged and His-tagged fusion proteins were expressed in BL21-Gold *E. coli* cells and purified as described in (38,57). GST-tagged proteins were bound to glutathione-agarose conjugated beads while His-tagged proteins were bound to Ni-NTA beads. Bound beads were then equilibrated in cleavage buffer (20 mM Tris pH 8.0, 150 mM NaCl, 2 mM EDTA, 10% glycerol and 1 mM DTT or BME) and incubated with tobacco etch virus protease overnight at 4°C to cleave proteins from the respective tag. Proteins were then purified by MonoQ or MonoS ion exchange chromatography at 4°C, followed by S200 gel filtration chromatography at 4°C. Purified proteins were eluted in 20 mM Tris pH 8.0, 150 mM NaCl, 10% glycerol, and 1 mM DTT, concentrated to working concentrations using Millipore column concentrator, and flash frozen in liquid nitrogen.

### Limited Proteolysis and Edman Degradation Sequencing

Proteins were diluted to 12μM in 20 mM Tris pH 8.0, 150 mM NaCl and 1 mM DTT and incubated at RT in 0.05 mg/mL sequencing grade trypsin (Roche Applied Science). Digestions were stopped with 2X Laemmli sample buffer and placed on ice until analysis. Samples were boiled and run by SDS-PAGE, then stained with 0.1% Coomassie Blue R-250, 40% ethanol, and 10% acetic acid. Gels were scanned on a LI-COR scanner. For N-terminal sequencing, digested peptides were blotted onto PVDF membrane, stained (0.1% Coomassie Blue R-250, 40% methanol, and 1% acetic acid), destained and dried. Individual bands were excised from the membrane and sequenced by Edman degradation (Iowa State University Protein Facility).

### Crosslinking Experiments

αT-catenin protein fragments were incubated with or without 1 mM BS3 (Thermo Scientific) in 20 mM HEPES, pH 7.4, 150 mM NaCl, and 1 mM DTT for 30 min at 4°C or 37°C, separated by SDS-PAGE, stained with Coomassie dye and imaged on a LI-COR scanner.

### Isothermal Titration Calorimetry

Proteins used for ITC were purified as described except the S200 buffer was replaced with ITC Buffer (20mM HEPES pH 8.0, 150mM NaCl, 1mM TCEP). Identical buffer was used to purify both cell and titrant samples to ensure buffer match. Only fresh, unfrozen proteins were used for ITC. Measurements were performed on a Malvern MicroCal PEAQ-ITC or MicroCal VP-ITC calorimeter (Malvern Panalytical). For experiments on the MicroCal PEAQ-ITC, titration occurred by an initial 0.5 μL injection followed by 18 x 2μL injections of 110-150 μM β-catenin, 103-158 μM β-catenin/N-cadherin tail complex or 396-600 μM vinculin aa 1-259 (D1) into the cell containing 9-55 μM of αT-catenin or αE-catenin. For experiments on the MircoCal VP-ITC, ligand was added with an initial 2 ul injection followed by 32 - 41 7 −9 ul injections. The concentration of αT-catenin head or αT-catenin head•β-catenin complex in the cell varied between 22 and 56 μM. Vinculin D1 concentrations in the syringe varied between 240 and 546 μM. All calorimetry experiments were performed at 25°C. All data analysis was performed using Malvern MicroCal ITC analysis software. For baseline correction, a mean baseline value, calculated by averaging the data points at saturation, was subtracted from each data point.

### Cell Culture

R2/7 carcinoma cells were cultured in DMEM (4.5 g/L glucose), 10% fetal bovine serum, 1mM sodium pyruvate and penicillin/streptomycin. Lipofectamine 2000 was used for all transient transfections.

### Immunostaining and Confocal Microscopy

Cells were fixed in 4% paraformaldehyde in PHEM buffer (60 mM 1,4-piperazin-ediethanesulfonic acid pH 7.0, 25 mM HEPES pH 7.0, 10 mM EGTA pH 8.0, 2 mM MgCl and 0.12 M sucrose), washed with PBS, permeabilized with 0.2% Triton X-100 in PBS for 2 minutes and then blocked for 1 hr at RT in PBS + 10% BSA. Samples were washed three times in PBS, incubated with primary in PBS + 1% BSA for 1 hr at RT, washed three times in PBS, incubated with secondary in PBS + 1% BSA for 1 hr at RT, washed three times in PBS, and mounted in ProLong Diamond mounting medium. Cells were imaged on a Nikon Eclipse Ti inverted microscope outfitted with a Prairie swept field confocal scanner, Agilent monolithic laser launch, and Andor iXon3 camera using NIS-Elements imaging software.

## Acknowledgements

We thank Jie Yan for sharing the titin I27-αT-catenin construct. We thank Cara Gottardi for the R2/7 cells.

## References

1. Ratheesh, A., and Yap, A. S. (2012) A bigger picture: classical cadherins and the dynamic actin cytoskeleton. Nature reviews. Molecular cell biology 13, 673–679

2. Rubsam, M., Broussard, J. A., Wickstrom, S. A., Nekrasova, O., Green, K. J., and Niessen, C. M. (2018) Adherens Junctions and Desmosomes Coordinate Mechanics and Signaling to Orchestrate Tissue Morphogenesis and Function: An Evolutionary Perspective. Cold Spring Harbor perspectives in biology 10

3. Mege, R. M., and Ishiyama, N. (2017) Integration of Cadherin Adhesion and Cytoskeleton at Adherens Junctions. Cold Spring Harbor perspectives in biology 9

4. Priest, A. V., Shafraz, O., and Sivasankar, S. (2017) Biophysical basis of cadherin mediated cellcell adhesion. Experimental Cell Research 358, 10–13

5. Shapiro, L., and Weis, W. I. (2009) Structure and biochemistry of cadherins and catenins. Cold Spring Harbor perspectives in biology 1, a003053

6. Meng, W. X., and Takeichi, M. (2009) Adherens Junction: Molecular Architecture and Regulation. Cold Spring Harbor perspectives in biology 1

7. Maiden, S. L., and Hardin, J. (2011) The secret life of alpha-catenin: moonlighting in morphogenesis. J Cell Biol 195, 543–552

8. Rimm, D. L., Koslov, E. R., Kebriaei, P., Cianci, C. D., and Morrow, J. S. (1995) Alpha 1(E)-catenin is an actin-binding and -bundling protein mediating the attachment of F-actin to the membrane adhesion complex. Proc Natl Acad Sci USA 92, 8813–8817

9. Yamada, S., Pokutta, S., Drees, F., Weis, W. I., and Nelson, W. J. (2005) Deconstructing the cadherin-catenin-actin complex. Cell 123, 889–901

10. Drees, F., Pokutta, S., Yamada, S., Nelson, W. J., and Weis, W. I. (2005) Alpha-catenin is a molecular switch that binds E-cadherin-beta-catenin and regulates actin-filament assembly. Cell 123, 903–915

11. Buckley, C. D., Tan, J., Anderson, K. L., Hanein, D., Volkmann, N., Weis, W. I., Nelson, W. J., and Dunn, A. R. (2014) Cell adhesion. The minimal cadherin-catenin complex binds to actin filaments under force. Science 346, 1254211

12. Pokutta, S., Choi, H. J., Ahlsen, G., Hansen, S. D., and Weis, W. I. (2014) Structural and Thermodynamic Characterization of Cadherin.beta-Catenin.alpha-Catenin Complex Formation. J Biol Chem 289, 13589–13601

13. Ishiyama, N., Sarpal, R., Wood, M. N., Barrick, S. K., Nishikawa, T., Hayashi, H., Kobb, A. B., Flozak, A. S., Yemelyanov, A., Fernandez-Gonzalez, R., Yonemura, S., Leckband, D. E., Gottardi, C. J., Tepass, U., and Ikura, M. (2018) Force-dependent allostery of the alpha-catenin actin-binding domain controls adherens junction dynamics and functions. Nat Commun 9, 5121

14. Ishiyama, N., Tanaka, N., Abe, K., Yang, Y. J., Abbas, Y. M., Umitsu, M., Nagar, B., Bueler, S. A., Rubinstein, J. L., Takeichi, M., and Ikura, M. (2013) An autoinhibited structure of alpha-catenin and its implications for vinculin recruitment to adherens junctions. J Biol Chem 288, 15913–15925

15. Rangarajan, E. S., and Izard, T. (2013) Dimer asymmetry defines alpha-catenin interactions. Nat Struct Mol Biol 20, 188–193

16. Terekhova, K., Pokutta, S., Kee, Y. S., Li, J., Tajkhorshid, E., Fuller, G., Dunn, A. R., and Weis, W. I. (2019) Binding partner- and force-promoted changes in alphaE-catenin conformation probed by native cysteine labeling. Sci Rep 9, 15375

17. Nieset, J. E., Redfield, A. R., Jin, F., Knudsen, K. A., Johnson, K. R., and Wheelock, M. J. (1997) Characterization of the interactions of alpha-catenin with alpha-actinin and beta-catenin/plakoglobin. J Cell Sci 110 (Pt 8), 1013–1022

18. Koslov, E. R., Maupin, P., Pradhan, D., Morrow, J. S., and Rimm, D. L. (1997) Alpha-catenin can form asymmetric homodimeric complexes and/or heterodimeric complexes with beta-catenin. J Biol Chem 272, 27301–27306

19. Pokutta, S., and Weis, W. I. (2000) Structure of the dimerization and beta-catenin-binding region of alpha-catenin. Mol Cell 5, 533–543

20. Imamura, Y., Itoh, M., Maeno, Y., Tsukita, S., and Nagafuchi, A. (1999) Functional domains of alpha-catenin required for the strong state of cadherin-based cell adhesion. J Cell Biol 144, 1311–1322

21. Pokutta, S., Drees, F., Takai, Y., Nelson, W. J., and Weis, W. I. (2002) Biochemical and structural definition of the l-afadin- and actin-binding sites of alpha-catenin. J Biol Chem 277, 18868–18874

22. Yang, J., Dokurno, P., Tonks, N. K., and Barford, D. (2001) Crystal structure of the M-fragment of alpha-catenin: implications for modulation of cell adhesion. EMBO J 20, 3645–3656

23. Choi, H. J., Pokutta, S., Cadwell, G. W., Bobkov, A. A., Bankston, L. A., Liddington, R. C., and Weis, W. I. (2012) alphaE-catenin is an autoinhibited molecule that coactivates vinculin. Proc Natl Acad Sci U S A 109, 8576–8581

24. Yonemura, S., Wada, Y., Watanabe, T., Nagafuchi, A., and Shibata, M. (2010) alpha-Catenin as a tension transducer that induces adherens junction development. Nat Cell Biol 12, 533–542

25. Li, J., Newhall, J., Ishiyama, N., Gottardi, C., Ikura, M., Leckband, D. E., and Tajkhorshid, E. (2015) Structural Determinants of the Mechanical Stability of alpha-Catenin. J Biol Chem 290, 18890–18903

26. Xu, X. P., Pokutta, S., Torres, M., Swift, M. F., Hanein, D., Volkmann, N., and Weis, W. I. (2020) Structural basis of alphaE-catenin-F-actin catch bond behavior. eLife 9

27. Mei, L., Espinosa de Los Reyes, S., Reynolds, M. J., Leicher, R., Liu, S., and Alushin, G. M. (2020) Molecular mechanism for direct actin force-sensing by alpha-catenin. eLife 9

28. le Duc, Q., Shi, Q., Blonk, I., Sonnenberg, A., Wang, N., Leckband, D., and de Rooij, J. (2010) Vinculin potentiates E-cadherin mechanosensing and is recruited to actin-anchored sites within adherens junctions in a myosin II,Äìdependent manner. The Journal of Cell Biology 189, 1107–1115

29. Yao, M., Qiu, W., Liu, R., Efremov, A. K., Cong, P., Seddiki, R., Payre, M., Lim, C. T., Ladoux, B., Mege, R. M., and Yan, J. (2014) Force-dependent conformational switch of alpha-catenin controls vinculin binding. Nat Commun 5, 4525

30. Barrick, S., Li, J., Kong, X. Y., Ray, A., Tajkhorshid, E., and Leckband, D. (2018) Salt bridges gate alpha-catenin activation at intercellular junctions. Molecular Biology of the Cell 29, 111–122

31. Maki, K., Han, S. W., Hirano, Y., Yonemura, S., Hakoshima, T., and Adachi, T. (2016) Mechano-adaptive sensory mechanism of alpha-catenin under tension. Sci Rep 6, 24878

32. Maki, K., Han, S. W., Hirano, Y., Yonemura, S., Hakoshima, T., and Adachi, T. (2018) Real-time TIRF observation of vinculin recruitment to stretched alpha-catenin by AFM. Sci Rep-Uk 8

33. Twiss, F., Le Duc, Q., Van Der Horst, S., Tabdili, H., Van Der Krogt, G., Wang, N., Rehmann, H., Huveneers, S., Leckband, D. E., and De Rooij, J. (2012) Vinculin-dependent Cadherin mechanosensing regulates efficient epithelial barrier formation. Biology open 1, 1128–1140

34. Merkel, C. D., Li, Y., Raza, Q., Stolz, D. B., and Kwiatkowski, A. V. (2019) Vinculin anchors contractile actin to the cardiomyocyte adherens junction. Mol Biol Cell, mbcE19040216

35. Thomas, W. A., Boscher, C., Chu, Y. S., Cuvelier, D., Martinez-Rico, C., Seddiki, R., Heysch, J., Ladoux, B., Thiery, J. P., Mege, R. M., and Dufour, S. (2013) alpha-Catenin and vinculin cooperate to promote high E-cadherin-based adhesion strength. J Biol Chem 288, 4957–4969

36. Zhao, Z. M., Reynolds, A. B., and Gaucher, E. A. (2011) The evolutionary history of the catenin gene family during metazoan evolution. BMC evolutionary biology 11, 198

37. Hulpiau, P., Gul, I. S., and van Roy, F. (2013) New Insights into the Evolution of Metazoan Cadherins and Catenins. Prog Mol Biol Transl 116, 71–94

38. Wickline, E. D., Dale, I. W., Merkel, C. D., Heier, J. A., Stolz, D. B., and Kwiatkowski, A. V. (2016) alphaT-Catenin Is a Constitutive Actin-binding alpha-Catenin That Directly Couples the Cadherin.Catenin Complex to Actin Filaments. J Biol Chem 291, 15687–15699

39. Janssens, B., Goossens, S., Staes, K., Gilbert, B., van Hengel, J., Colpaert, C., Bruyneel, E., Mareel, M., and van Roy, F. (2001) alphaT-catenin: a novel tissue-specific beta-catenin-binding protein mediating strong cell-cell adhesion. J Cell Sci 114, 3177–3188

40. Goossens, S., Janssens, B., Bonne, S., De Rycke, R., Braet, F., van Hengel, J., and van Roy, F. (2007) A unique and specific interaction between alphaT-catenin and plakophilin-2 in the area composita, the mixed-type junctional structure of cardiac intercalated discs. J Cell Sci 120, 2126–2136

41. Vite, A., and Radice, G. L. (2014) N-cadherin/catenin complex as a master regulator of intercalated disc function. Cell communication & adhesion 21, 169–179

42. Li, J., Goossens, S., van Hengel, J., Gao, E., Cheng, L., Tyberghein, K., Shang, X., De Rycke, R., van Roy, F., and Radice, G. L. (2012) Loss of alphaT-catenin alters the hybrid adhering junctions in the heart and leads to dilated cardiomyopathy and ventricular arrhythmia following acute ischemia. J Cell Sci 125, 1058–1067

43. van Hengel, J., Calore, M., Bauce, B., Dazzo, E., Mazzotti, E., De Bortoli, M., Lorenzon, A., Li Mura, I. E., Beffagna, G., Rigato, I., Vleeschouwers, M., Tyberghein, K., Hulpiau, P., van Hamme, E., Zaglia, T., Corrado, D., Basso, C., Thiene, G., Daliento, L., Nava, A., van Roy, F., and Rampazzo, A. (2013) Mutations in the area composita protein alphaT-catenin are associated with arrhythmogenic right ventricular cardiomyopathy. European heart journal 34, 201–210

44. Folmsbee, S. S., Morales-Nebreda, L., Van Hengel, J., Tyberghein, K., Van Roy, F., Budinger, G. R., Bryce, P. J., and Gottardi, C. J. (2015) The cardiac protein alphaT-catenin contributes to chemical-induced asthma. Am J Physiol Lung Cell Mol Physiol 308, L253–258

45. Folmsbee, S. S., Wilcox, D. R., Tyberghein, K., De Bleser, P., Tourtellotte, W. G., van Hengel, J., van Roy, F., and Gottardi, C. J. (2016) alphaT-catenin in restricted brain cell types and its potential connection to autism. J Mol Psychiatry 4, 2

46. Chiarella, S. E., Rabin, E. E., Ostilla, L. A., Flozak, A. S., and Gottardi, C. J. (2018) alphaT-catenin: A developmentally dispensable, disease-linked member of the alpha-catenin family. Tissue Barriers 6, e1463896

47. Pang, S. M., Le, S., Kwiatkowski, A. V., and Yan, J. (2019) Mechanical stability of alphaT-catenin and its activation by force for vinculin binding. Mol Biol Cell 30, 1930–1937

48. Kostetskii, I., Li, J., Xiong, Y., Zhou, R., Ferrari, V. A., Patel, V. V., Molkentin, J. D., and Radice, G. L. (2005) Induced deletion of the N-cadherin gene in the heart leads to dissolution of the intercalated disc structure. Circulation research 96, 346–354

49. Watabe-Uchida, M., Uchida, N., Imamura, Y., Nagafuchi, A., Fujimoto, K., Uemura, T., Vermeulen, S., van Roy, F., Adamson, E. D., and Takeichi, M. (1998) alpha-Catenin-vinculin interaction functions to organize the apical junctional complex in epithelial cells. J Cell Biol 142, 847–857

50. Li, Y., Merkel, C. D., Zeng, X., Heier, J. A., Cantrell, P. S., Sun, M., Stolz, D. B., Watkins, S. C., Yates, N. A., and Kwiatkowski, A. V. (2019) The N-cadherin interactome in primary cardiomyocytes as defined by quantitative proximity proteomics. J Cell Sci

51. Sheikh, F., Chen, Y., Liang, X., Hirschy, A., Stenbit, A. E., Gu, Y., Dalton, N. D., Yajima, T., Lu, Y., Knowlton, K. U., Peterson, K. L., Perriard, J. C., and Chen, J. (2006) alpha-E-catenin inactivation disrupts the cardiomyocyte adherens junction, resulting in cardiomyopathy and susceptibility to wall rupture. Circulation 114, 1046–1055

52. Kowalczyk, A. P., and Green, K. J. (2013) Structure, function, and regulation of desmosomes. Progress in molecular biology and translational science 116, 95–118

53. Choi, H. J., Gross, J. C., Pokutta, S., and Weis, W. I. (2009) Interactions of plakoglobin and beta-catenin with desmosomal cadherins: basis of selective exclusion of alpha- and beta-catenin from desmosomes. J Biol Chem 284, 31776–31788

54. Benjamin, J. M., Kwiatkowski, A. V., Yang, C., Korobova, F., Pokutta, S., Svitkina, T., Weis, W. I., and Nelson, W. J. (2010) AlphaE-catenin regulates actin dynamics independently of cadherin-mediated cell-cell adhesion. J Cell Biol 189, 339–352

55. Wood, M. N., Ishiyama, N., Singaram, I., Chung, C. M., Flozak, A. S., Yemelyanov, A., Ikura, M., Cho, W., and Gottardi, C. J. (2017) alpha-Catenin homodimers are recruited to phosphoinositide-activated membranes to promote adhesion. J Cell Biol 216, 3767–3783

56. Huveneers, S., Oldenburg, J., Spanjaard, E., van der Krogt, G., Grigoriev, I., Akhmanova, A., Rehmann, H., and de Rooij, J. (2012) Vinculin associates with endothelial VE-cadherin junctions to control force-dependent remodeling. J Cell Biol 196, 641–652

57. Hansen, S. D., Kwiatkowski, A. V., Ouyang, C. Y., Liu, H., Pokutta, S., Watkins, S. C., Volkmann, N., Hanein, D., Weis, W. I., Mullins, R. D., and Nelson, W. J. (2013) alphaE-catenin actin-binding domain alters actin filament conformation and regulates binding of nucleation and disassembly factors. Mol Biol Cell 24, 3710–3720

